# Histology to MRI registration quality of *ex vivo* human brain blocks fixed with solutions used in anatomy laboratories

**DOI:** 10.1101/2025.05.06.652445

**Authors:** Eve-Marie Frigon, Amy Gérin-Lajoie, Jérémie P. Fouquet, Yashar Zeighami, Denis Boire, Mahsa Dadar, Josefina Maranzano

## Abstract

**Introduction:** *Post-mortem* brain tissue is obtained from brain banks that provide small tissue samples, while gross anatomy laboratories could become a source of complete brains for neuroscientists. These are preserved with solutions whose chemical composition differs from the classic neutral-buffered formalin (NBF) used in brain banks, such as a saturated-salt-solution (SSS) or an alcohol-formaldehyde solution (AFS) that preserve antigenicity of the main brain cell populations. Since histology remains the gold standard in neuroscientific research, while MRI is the most common imaging modality, MRI-histology registration quality needs to be assessed to ensure the suitability using brains fixed with innovative solutions for research procedures. Hence, our goal was to compare the registration quality of human brains fixed with NBF, SSS and AFS, as well as the histological characteristics that could affect the registration.

**Methods:** We used 12 human brain blocks of 3×3×3 cm^3^ fixed in our anatomy laboratory using SSS (N=4), AFS (N=4), or NBF (N=4). The blocks were scanned using a 7Tesla Bruker animal MRI scanner with a T2-TurboRARE sequence at 0.13×0.13×0.5mm^3^. Blocks were then cut into 40μm thick sections (parallel to the 0.13×0.13 plane) using a vibratome. Sections were stained with histochemistry (HC) (Cresyl violet, Prussian blue, Luxol fast blue, H&E, and Bielschowsky) and immunohistochemistry (IHC) of the 4 main cell populations: neurons (NeuN), astrocytes (GFAP), microglia (Iba1) and myelin (PLP) either with or without an antigen retrieval (AR) protocol. Stained sections were imaged using a slide scanner microscope and segmented with masks using Display, and the sections were manually registered to the T2-TurboRare images using landmarks in Register (MincToolKit).

**Results:** More landmarks were needed to achieve proper alignment of the histology to MRI images for the SSS-fixed blocks, due to the lower GM-WM contrast in these brains. However, there was no significant difference in the staining intensity of the histology sections of blocks fixed with the three solutions, while SSS-fixed blocks showed a lower percentage of overlap between the good histology quality masks and the MRI masks. This resulted in a sufficient registration quality of all blocks, although more challenging when fixed with SSS.

**Conclusion:** We have developed histology, MRI and registration protocols that are of good quality in brain blocks fixed with solutions used in gross anatomy laboratories. These results are promising for neuroscientists interested in using full brains from anatomy laboratories, either using MRI, histology or registration of both modalities to study normal aging and neurodegenerative conditions.

## 1. Introduction

Magnetic resonance imaging (MRI) is the most common neuroimaging modality in neuroscientific research, since it is non-invasive and can provide good contrast between the cortical gray matter (GMc) and white matter (WM), enabling assessment and quantification of different brain structures and pathologies in a non-invasive manner. Different MRI acquisition protocols and technologies have been developed in the last few decades, increasing our ability to estimate whole brain, GMc and WM volumes,^1^ volumetric and relaxation changes of specific structures in neurodegenerative diseases (e.g., hippocampal volume changes in dementia),^2,3^ and lesion occurrence and their longitudinal evolution (e.g., white matter hyperintensities in cerebrovascular disease).^4^ However, to confirm MRI findings, histology remains the gold standard in neuroscientific research since it provides cellular resolution.^5,6^ Being able to correlate MRI imaging with cellular histology is of utmost importance to establish a meaningful anatomical and pathophysiological conclusion.

Biopsies from living individuals for histopathological study can only provide very small samples and are rarely performed due to their invasive nature. Therefore, researchers commonly use *post-mortem* tissue samples that are generally provided by brain banks. However, brain banks can usually only provide limited numbers of small tissue samples, one hemisphere being fixed by immersion in Neutral-buffered formalin (NBF), and the other frozen.^7,8^ Alternatively, neuroscientists could obtain higher numbers of full brains from gross anatomy laboratories that collaborate with body donation programs. These brains are fixed through whole-body perfusion, using solutions other than NBF typically used in brain banks since the body is used for teaching gross anatomy. Previous work of our team has shown that two solutions used in our human anatomy laboratory, either a saturated-salt solution (SSS) and an alcohol-formaldehyde solution (AFS), preserve antigenicity of the four main cell populations in mice brains^9^ and in human brain samples,^10^ potentially increasing the amount of brain tissue available to neuroscientists for histological studies. Furthermore, performing MRI scans in these brains is also feasible, enabling correlations with histology findings.^11,12^

Since MRI is the most common neuroimaging modality, and histology remains the gold standard, registration of images obtained with these two modalities is essential to correlate findings.^13-15^ Previous work on Alzheimer’s disease,^16-21^ Parkinson’s disease,^22^ strokes,^23^ cerebrovascular diseases,^24-27^ white matter hyperintensities (WMH),^28,29^ multiple sclerosis,^30-33^ etc., have used registration between MRI and histology to either confirm the clinical diagnosis or to elucidate the cellular changes in tissues and lesions that are characteristic of these pathologies.

However, while both MRI and histology assessments are feasible in tissues fixed by perfusion using neuroanatomy solutions, previous work has not established whether studies that use registration would be feasible and of sufficient quality in these brains. Hence, the goal of the present study was to describe the protocols that are suitable for linear registration of the histology sections to MRI in human brain blocks fixed with three solutions (SSS and AFS used in anatomy laboratories, and NBF used in brain banks), and to compare the robustness of the alignments in different histology stains across the fixatives.

## 2. Methods

### 2.1 Population

We used a convenience sample of 12 human brain blocks (SSS: N=4, AFS: N=4, and NBF: N=4) from our body donation program (Université du Québec à Trois-Rivières), that were fixed by whole-body perfusion. Prior to death, the donors consented to body donation and sharing their medical information for research or teaching purposes. This study was approved by the University’s Ethic Subcommittee on Anatomical Teaching and Research (Université du Québec à Trois-Rivières) and by the Douglas Research Centre Ethic Committee (McGill University, Montreal). The mean age at time of death was 81 years old (SD=9.65; range from 64 to 96). Female to male ratio reached 2:1. The mean *post-mortem* interval (PMI, i.e., delay between death and fixation of the donor) was 18.6 hours (SD=8.6; range from 3 to 36). The donors’ demographics are reported in Table 1.

**Table 1.** Characteristics of the specimens included in this study.

| Fixative | Sex | Age | Cause of death | PMI (hours) | Lobe |
| --- | --- | --- | --- | --- | --- |
| NBF | M | 74 | Lung cancer | 3 | Frontal R |
|  |  | 85 | Pulmonary oedema | 12 | Parietal R |
|  | F | 84 | Sepsis | 21 | Temporal L |
|  |  | 89 | Pneumonia | 23 | Occipital R |
| SSS | M | 83 | Cardiac arrest | 20 | Occipital R |
|  | F | 67 | COPD | 36 | Parietal L |
|  |  | 75 | Pneumonia and MS | 27 | Temporal R |
|  |  | 87 | Pneumonia | 9.5 | Frontal L |
| AFS | M | 91 | COPD | 20 | Parietal L |
|  | F | 64 | Pneumonia, dementia | 15.5 | Occipital R |
|  |  | 96 | Breast cancer | 13.5 | Temporal L |
|  |  | 80 | Cancer | 23 | Frontal L |

NBF=Neutral-buffered formalin; SSS=Saturated-salt solution; AFS=Alcohol-formaldehyde solution; F=Female; M=Male; COPD=Chronic obstructive pulmonary disease; MS=Multiple Sclerosis; PMI=*post-mortem* interval.

### 2.2 Magnetic resonance imaging protocol

Tissue blocks of approximately 3×3×3 cm^3^ were cut from regions that were macroscopically of good quality, which included: 1-a good color (beige surface without remaining blood); 2-a firm texture, and 3-free of tears that could have happened during brain extraction or removal of the arachnoid. Blocks were packed in small containers filled with a high-density fluoropolymer fluid free of MRI signal (Christo-Lube, Engineered Custom Lubricants). To avoid air bubbles, the tissue blocks were packed in the container while fully immersed in a bucket filled with the fluoropolymer fluid, closing the lid while the specimen and container were fully immersed. Blocks were then scanned in a 7 Tesla Bruker Biospec 70/30 animal MRI scanner at the Douglas Cerebral Imaging Centre (CIC) using a Bruker transmit-receive volume coil with an 86-mm inner diameter. Six high-resolution sequences (T1-FLASH, T2-TurboRARE, T1-map, T2-map, T2*map, and MP2RAGE) were acquired overnight using parameters that were described in the Douglas-Bell Canada Brain Bank Post-mortem brain imaging protocol.^34^

### 2.3 Histology procedures

#### 2.3.1 Sectioning and mounting

After MRI scans were completed, the tissue blocks were rinsed in 0.1M Phosphate buffer and photographed. The blocks that were too large to fit into the vibratome (Leica VT1000S) were cut and photographed again before cutting into 40 µm sections. Sections of two different levels of the blocks (also photographed), parallel to the high MRI resolution plane, were selected as follows: five sections per level (ten total) were mounted on 70 × 50 mm slides and processed with five histochemistry (HC) stains. Similarly, two sections from the same level were mounted for immunohistochemistry (IHC) with 4 different antibodies. We selected the level that showed the least amount of deformation and tears to obtain these eight sections. One section per antibody was processed with a heat-induced antigen retrieval (AR) protocol (boiling 20 minutes in 0.01M Citrate Buffer), and the other was processed with IHC directly, to avoid the potential additional tearing and deformation caused by the heat of the AR procedure. This procedure resulted in 18 mounted sections per block.

#### 2.3.2 Immunohistochemistry

IHC was used to label the four main cell populations: neurons, using neuronal nuclei (NeuN) antibodies, astrocytes, using glial fibrillary acidic protein (GFAP) antibody, microglia, using ionized binding adaptor molecule 1 (Iba1) antibody and oligodendrocytic myelin using proteolipid protein (PLP) antibody. Procedures were followed as described in Frigon et al., 2024^10^ but sections were processed mounted on the slides rather than free-floating.

#### 2.3.3 Histochemistry stains

We used five HC stains to label the main brain cell populations, avoiding antibodies that require antigen configuration specificity. We used Cresyl Violet (Nissl stain, which labels neurons and glial cell bodies), Luxol Fast Blue (myelin), Prussian blue staining (Perl’s stain, which labels iron deposits), Bielschowsky’s silver staining (which labels axons and neurofibrillary tangles) as previously described in Frigon et al., 2024.^10^ We also used Hematoxylin and Eosin (H&E) Staining Kit from Abcam (ab245880) and followed their procedures.

#### 2.3.4 Imaging

Mounted stained sections were scanned at 40X magnification with an Olympus VS120 Slide scanner at the CIC. The images were converted from .vsi format to .nifti format and downsampled by a factor of 20 to reduce file size to facilitate the registration step using an in-house MATLAB script. The resulting images were converted to minc format to allow the use of the Register software from the MINC Tool-Kit (McConnell Brain Imaging Centre) to perform manual registration.

### 2.4 Registration

The T2-TurboRARE images (voxel resolution of 0.13*0.13*0.5 mm) were used for registration with histology since they provide good anatomical contrast that are close to *in vivo* images, in comparison to *post-mortem* T1w images in which the gray level contrast is reversed.^34^ Registration between histology images and MRIs was performed using a manual linear registration approach (3 rotations, 3 translations, 1 scale), tagging a minimum of 4 landmarks using Register software (included in the MINC Tool-Kit). More fiducial markers (i.e., tags) were placed if necessary, and the required number was considered as a variable of interest (see next section). Since all the acquired *post-mortem* sequences are co-registered together, histology images could also be co-registered to the other acquired sequences using the same transformations. The same approach was repeated to register the block photos to the MRIs.

### 2.5 Variables of interest

#### 2.5.1 Histology quality

##### 2.5.1.1 Staining intensity

Histological staining intensity of the sections was qualitatively assessed as it could impact the alignment of the tissue, i.e., if certain parts of the tissue are not sufficiently stained, then those areas could be impossible to align. A darker section reflected the better quality of the histology.

Pale=0 (Figure 1A); Heterogeneous=1 (Figure 1B); Dark=2 (Figure 1C).

**Figure 1.**
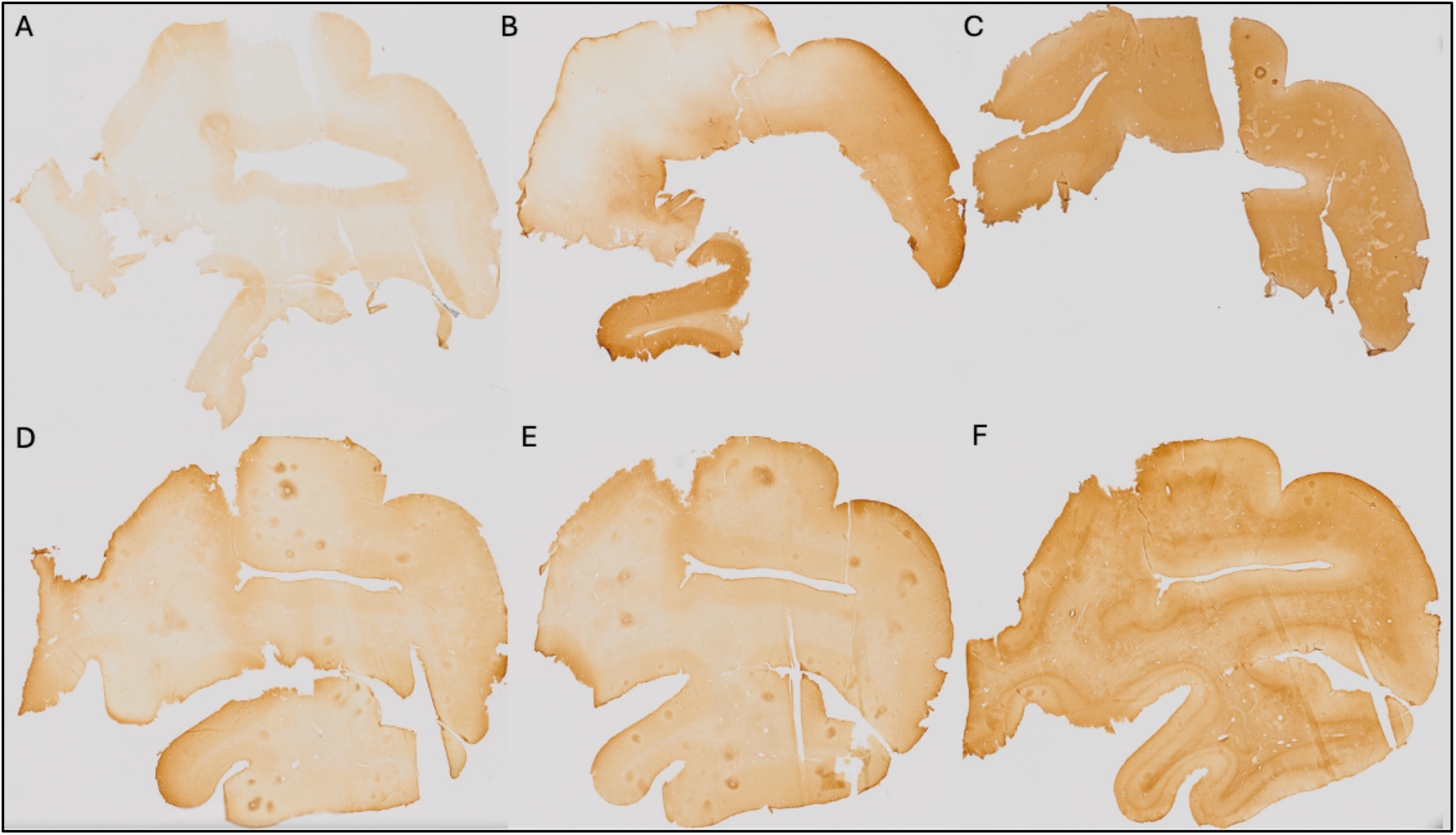
Criteria for assessment of the histology quality variable. Staining intensity showing A) Pale section=0; B) Heterogeneous section=1 and C) Dark section=2. GM-WM contrast showing D) No contrast=0; E) Blurred contrast=1 and F) Sharp contrast=2.

##### 2.5.1.2 Gray matter to white matter contrast

The GM-WM contrast was assessed since more contrast on the histology sections allows for better placement of markers on different boundaries of the tissue (i.e., pial surface of the GMc and GMc-to-WM boundary). A sharp GM-WM contrast reflected a better quality of the histology.

No contrast=0 (Figure 1D); Blurred contrast=1 (Figure 1E); Sharp contrast=2 (Figure 1F).

#### 2.5.2 Registration quality

##### 2.5.2.1 Number of tags

The number of tags that were used for manual registration was considered as a variable that could inform on the ease of registration. If more than 4 tags were needed (minimum to initiate manual alignment), this could reflect that the registration was more complicated. An example of the anatomical landmarks (6 tags) that were used is shown in Figure 2.

**Figure 2.**
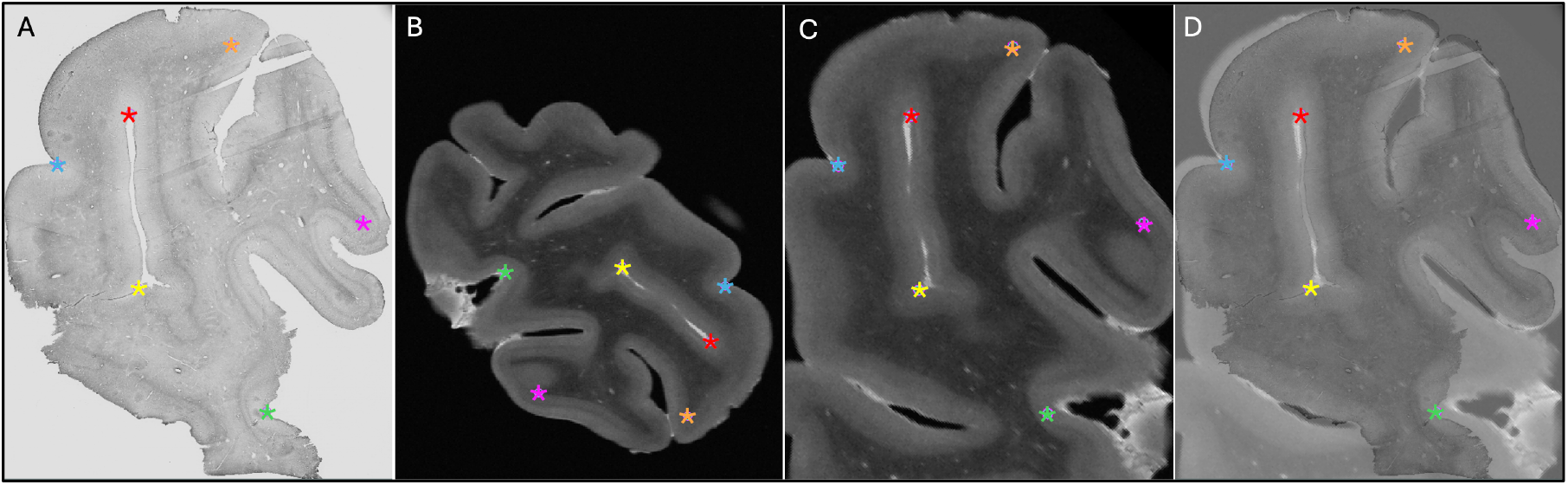
Process of the manual registration using landmarks. A) Histology section. B) Native T2-TurboRARE MRI sequence. C) T2-TurboRARE MRI registered to the histology section. D) Overlap of the histology and registered MRI. Colored stars (6 tags in total: orange, red, blue, pink, yellow and green) show the landmarks that were used to initiate the registration between A and B, which resulted in C, and the overlap of both modalities in D.

##### 2.5.2.2 Tissue block-MRI-histology colocalization

Using Display software from the MINC toolkit, we segmented the surfaces of the same level of the MRI slide (Figure 3A) and the photograph of the tissue block (Figure 3B) registered to the corresponding histology section (Figure 3C). The histology sections were segmented by an expert with more than 8 years of experience (EMF), creating the following masks: Label 1 (Red)=Non-folded/Non-torn Tissue (Good quality); Label 2 (Green)=Folded Tissue; Label 3 (Blue)=Torn Tissue; and Label 4 (Cyan)=Non-folded/Non-torn Tissue that was not parallel to the MRI plane (Good quality but off plane) (Figure 3C). The surface of each mask was calculated as number of pixels in the MRI image space by transforming the histology and tissue block images to the MRI image space using the generation transforms. The MRI and photo block masks (Figure 3A-B) were then overlapped (Figure 4A) to assess the amount of tissue that was cut from the scanned block to be able to fit the vibratome. Then, the overlap of the histology mask (Label 1, Non-folded/Non-Teared Tissue) and the photo block mask (Figure 3B-C and Figure 4B) was used to assess the percentage of tissue loss in the histology sections. Dice Kappas of the overlapped masks were also calculated to assess the amount of overlap. All measurements were generated automatically using an in-house MATLAB script. The percentage of tissue showing Good quality was then compared according to the fixatives and the staining types. We also assessed the mean percentage of Folded Tissue and Torn Tissue for each group to assess if the quality of the histology section affected the registration quality.

**Figure 3.**
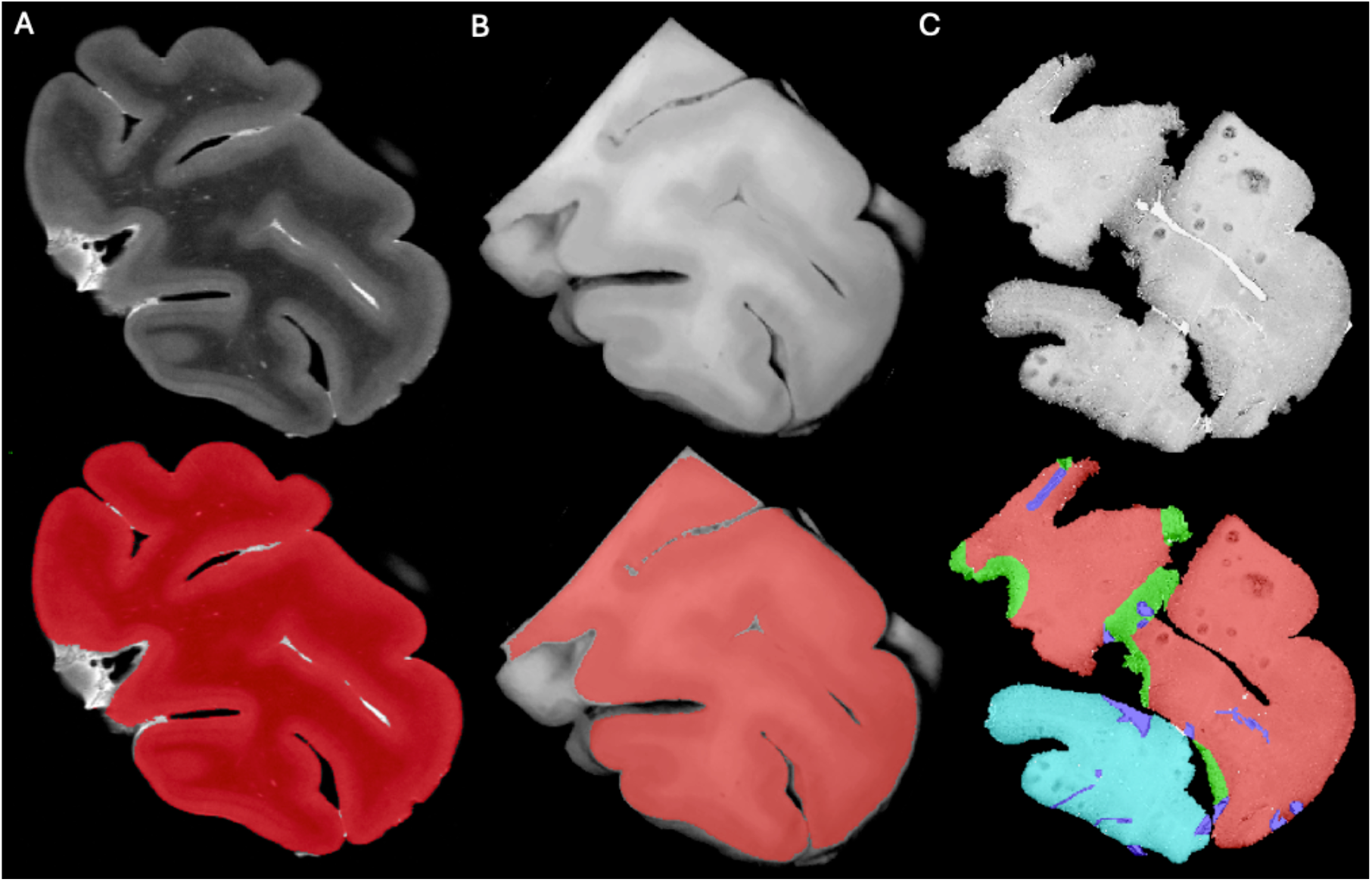
Masks created using Display to assess histology-MRI colocalization. Masks were created for A) MRI slide registered to the histology section; B) Photo of the block cut to fit in the vibratome, registered to the MRI-histology section; C) Histology sections photomicrographs. Red masks=Non-folded/Non-torn Tissue (Good quality); Green masks=Folded Tissue; Blue masks=Torn Tissue; and Cyan blue= Non-folded/Non-torn Tissue that was not parallel to the MRI plane (Good quality but off plane).

**Figure 4.**
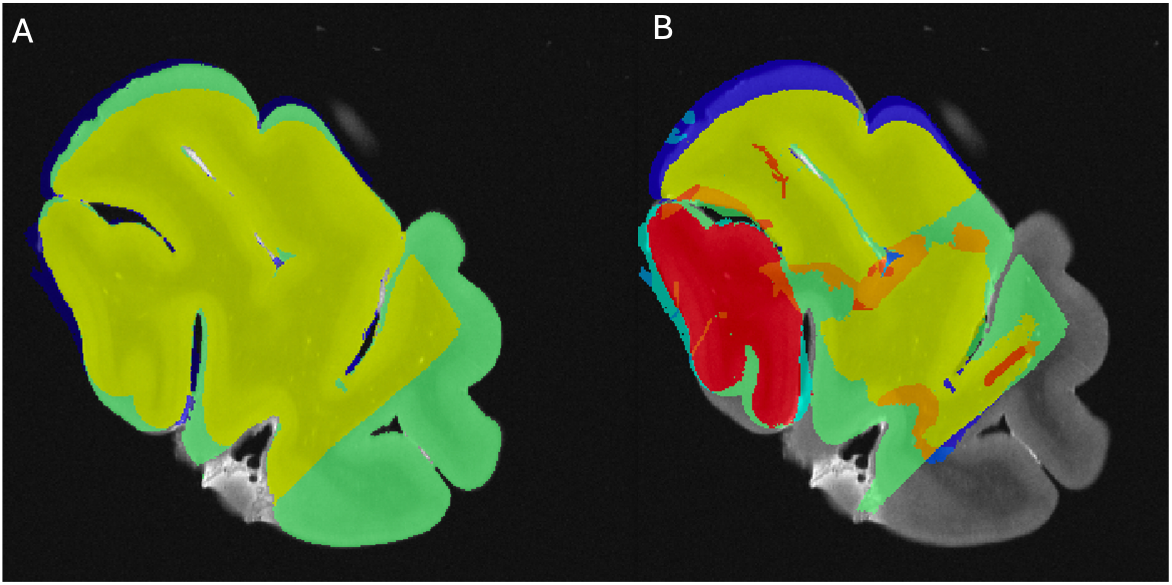
Overlap of the masks segmented to assess the amount of histology tissue colocalizing with MRI scans. A) Overlap of the MRI plane and block photo masks that was considered as the target for assessment of the histology sections that co-localized. B) Overlap of the histology (with the four labels) to the block photo masks that is overlapped with the MRI picture, which was assessed according to the two groups (fixatives and type of staining).

### 2.6 Statistical analysis

Qualitative variables (section 2.5.1: staining intensity and GM-WM contrast) were assessed using chi-square. Quantitative variables (section 2.5.2: number of tags and tissue block-MRI-histology colocalization) were assessed using Kruskal-Wallis analyses according to their distribution. All the variables were compared between the three experimental groups (NBF, SSS and AFS), as well as between the three different staining types (IHC with AR, IHC without AR and HC). All statistical analyses were conducted using IBM SPSS statistics (version 29.0.2.0) and were adjusted for multiple comparisons following a Bonferroni correction.

## 3. Results

### 3.1 Histology quality

#### 3.1.1 Staining intensity

No significant difference was found in the staining intensity of the histology sections across the three fixatives (p=0.696) (Figure 5, top left). IHC with AR showed more sections with a heterogeneous staining intensity, but this did not retain significance after a Bonferroni correction (uncorrected p=0.04). IHC without AR showed a significantly higher number of Pale sections than with AR and HC staining (p<0.0001) and significantly fewer Dark sections (p=0.0003). Finally, HC sections showed significantly fewer Pale sections (p=0.0007) and a significantly higher number of Dark sections (p<0.0001) (Figure 5, top right).

**Figure 5.**
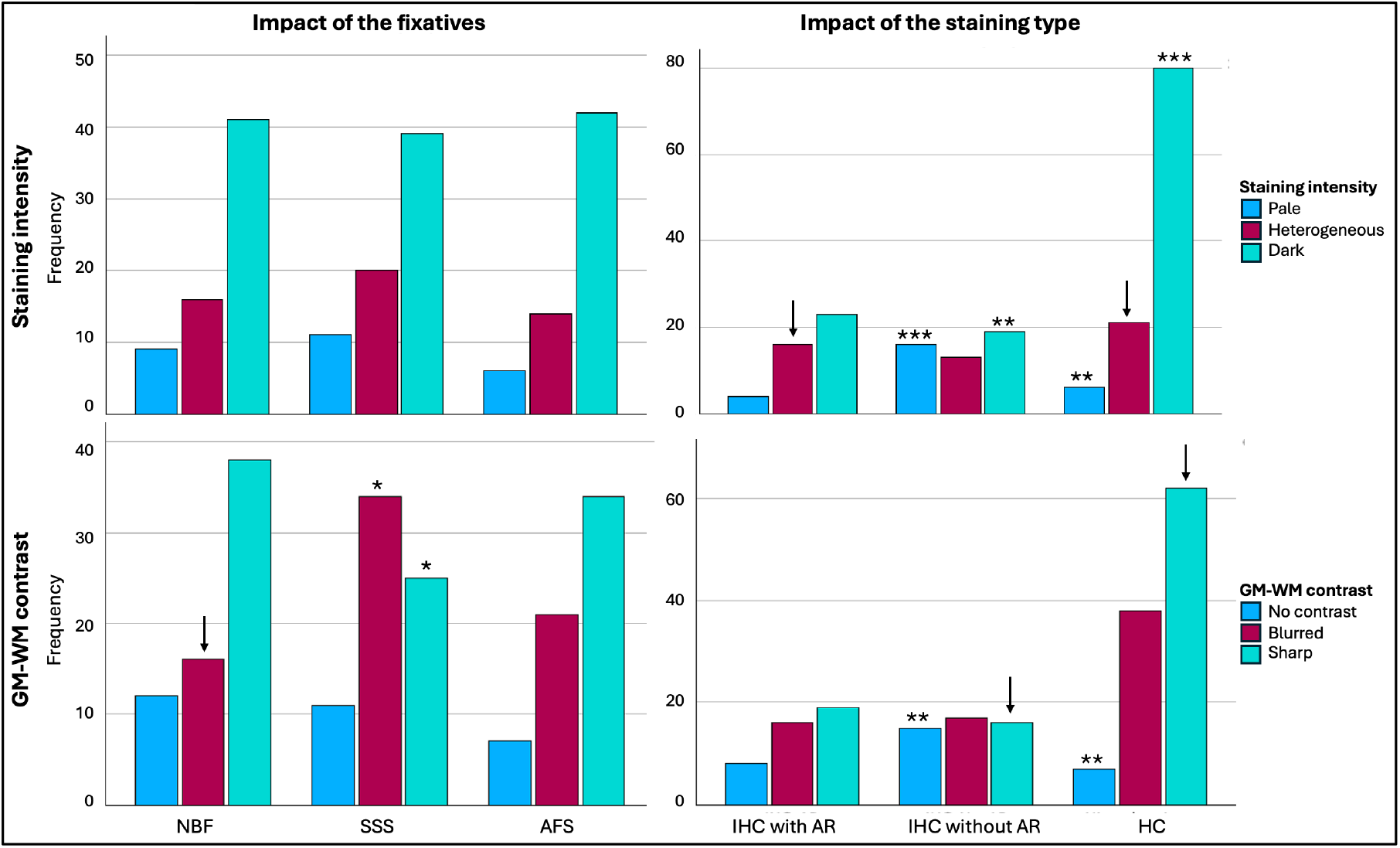
Histology quality of the sections regarding staining intensity and gray to white matter (GM-WM) contrast. Left graphs show the impact of the fixative (NBF=Neutral-buffered formalin; SSS=Saturated-salt solution; AFS=Alcohol-formaldehyde solution) on the staining intensity (top) and GM-WM contrast (bottom). Right graphs show the impact of the staining type (IHC=Immunohistochemistry; AR=Antigen retrieval; HC= Histochemistry) on the staining intensity (top) and GM-WM contrast (bottom). ↓p<0.05 before Bonferroni correction; *p<0.005; **p<0.001; ***p<0.0001.

#### 3.1.2 Gray matter to white matter contrast

GM-WM contrast showed significant differences between sections of blocks that were fixed with the three solutions (p=0.03). NBF-fixed blocks showed fewer blurred sections than the other two solutions, but this did not retain significance after a Bonferroni correction (uncorrected p=0.016). SSS-fixed blocks showed a significantly higher number of Blurred contrast (p=0.005) and significantly fewer Sharp contrast (p=0.005) than the sections fixed with the other two solutions (Figure 5, bottom left). GM-WM contrast was also significantly affected by the staining type (p=0.001). IHC with AR did not show any statistical difference. However, IHC without AR showed significantly more sections with No contrast (p=0.0003) and fewer sections with a Sharp contrast than HC, but this did not retain significance after a Bonferroni correction (uncorrected p=0.012). Finally, HC staining showed significantly fewer sections with No contrast (p=0.0002) and more sections with Sharp contrast, but this did not retain significance after Bonferroni correction (uncorrected p=0.0069) (Figure 5, bottom right).

### 3.2 Registration quality

#### 3.2.1 Number of tags

SSS-fixed blocks required significantly more tags to initiate alignment (median=6, range 4-16) than the blocks that were fixed with the other two solutions (NBF; median=5, range 4-25; p=0.002 and AFS, median=4, range 4-10; p<0.001) (Figure 6A). The staining type (IHC with AR, IHC without AR and HC) did not have an impact on the number of tags that were necessary to manually register the sections, since no significant difference (p=0.55) was found across groups (IHC-AR, median=6, range 4-25; IHC-noAR, median=5, range 4-9; HC, median=5, range 4-16) (Figure 6B).

**Figure 6.**
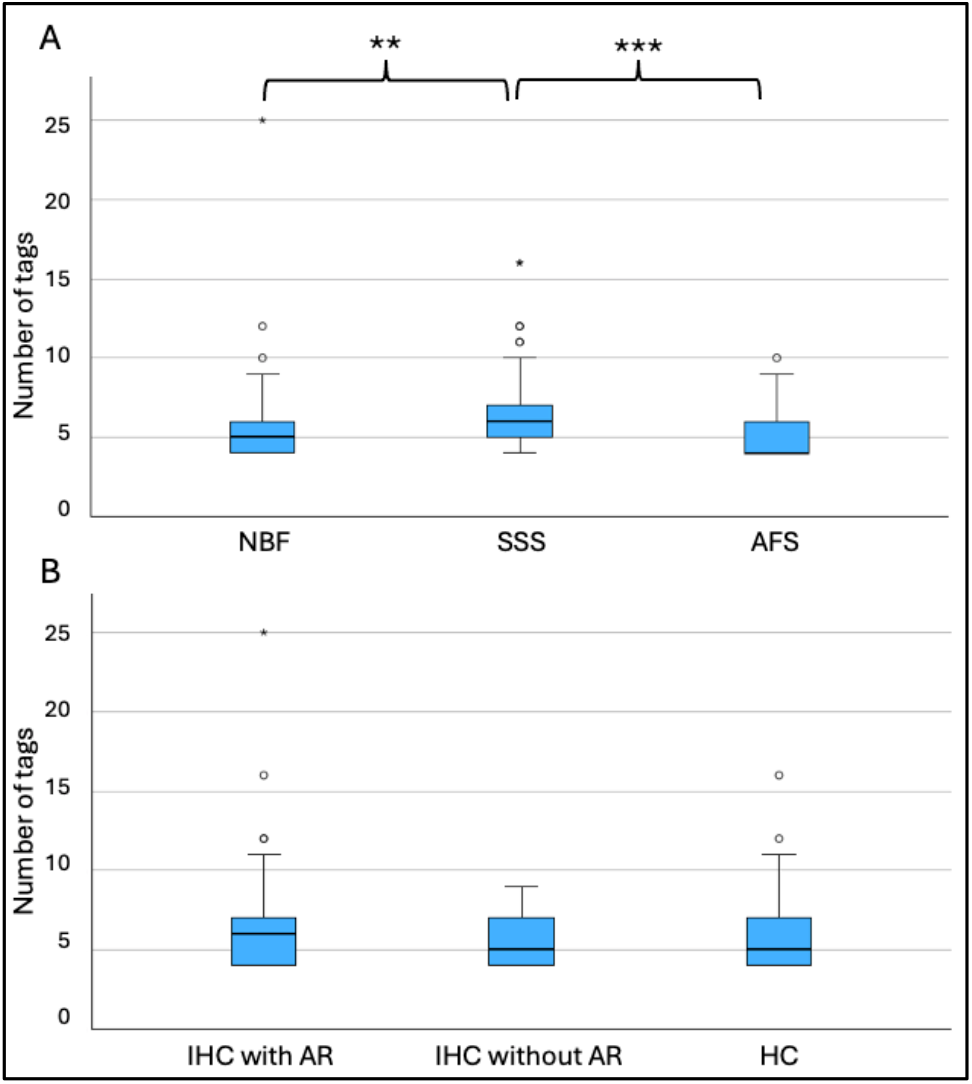
Number of tags necessary to register the sections. A) according to the fixative and B) according to the type of staining. NBF=Neutral-buffered formalin SSS=Saturated-salt solution; AFS=Alcohol-formaldehyde solution; IHC=Immunohistochemistry; AR=Antigen retrieval; HC=Histochemistry. **p<0.001; ***p<0.0001.

#### 3.2.2 Tissue block-MRI-histology colocalization

We used the overlap of the photo block and MRI (Figure 4A) as the target of the histology masks to assess the percentage of the histology sections of Good quality (label 1=Non-folded/Non-torn tissue) that colocalized with the MRIs. AFS-fixed blocks showed significantly higher overlap (%) between the MRI and Good quality histology mask (median=80.49, range 18-97, median kappa=0.892) than SSS-fixed blocks (median=67.40, range 0-93, median kappa=0.805) (p=0.012) (Figure 7 top left). Percentage of the good quality histology masks overlap with MRI and tissue block overlap were significantly different between the staining types, with a median of IHC with AR, IHC without AR and HC of 50.78 (range 0-96, median kappa=0.674), 76.45 (range 25-97, median kappa=0.841) and 72.46 (range 18-96, median kappa=0.870) respectively (p<0.001). IHC with AR histology sections showed significantly less overlap than IHC without AR (p<0.001) and HC (p=0.003) (Figure 7 top right).

**Figure 7.**
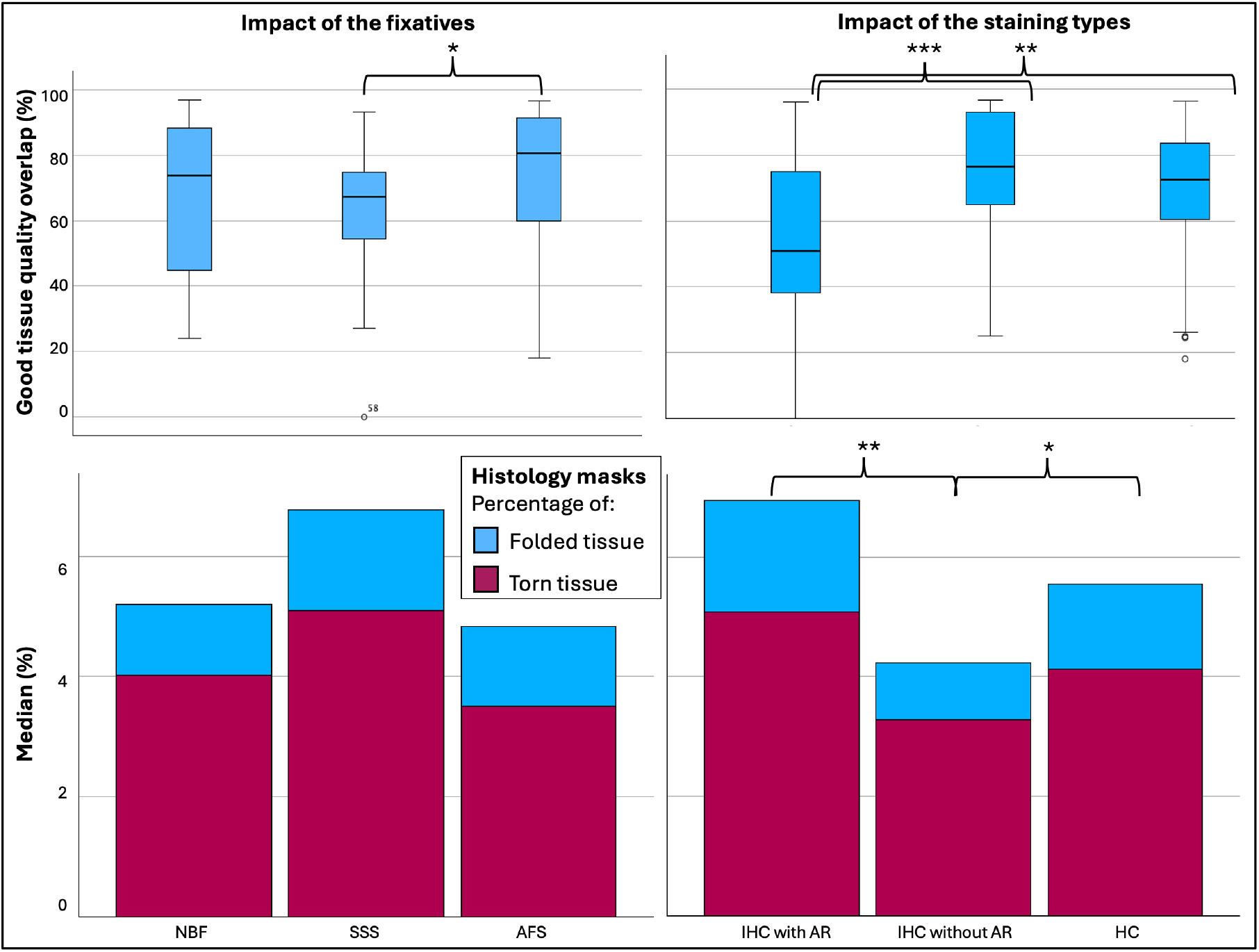
Tissue block-MRI-histology overlap of the masks to assess colocalization. Left graphs show the impact of the fixative (NBF=Neutral-buffered formalin; SSS=Saturated salt solution; AFS=Alcohol-formaldehyde solution) on the percentage of the Good quality tissue overlap (top) and the Median percentage of Folded (blue) and Torn (purple) tissue (bottom). Right graphs show the impact of the staining type (IHC=Immunohistochemistry; AR=Antigen retrieval; HC= Histochemistry) on the percentage of the Good quality tissue overlap (top) and the Median percentage of Folded (blue) and Torn (purple) tissue (bottom). *p<0.05; **p<0.01; ***p<0.001.

We also found that there was a higher percentage of Torn tissue in the brain blocks fixed with SSS, but this did not retain significance after a Bonferroni correction (Figure 7 bottom left). The histology sections processed with IHC without AR (median=3.24, range 0.82-23.60) showed significantly less Folded tissue than sections processed with HC (median=4.13, range 0-25.57, p=0.027) and with IHC with AR (median=5.08, range=1.35-71.40, p=0.004) (Figure 7 bottom right).

## 4. Discussion

This study aimed to describe the quality of linear registration between histology sections and MRI in human brain blocks fixed with three solutions, and in histology sections that were processed with different staining types to provide new insights on registration protocols of brain blocks from gross anatomy laboratories.

### 4.1 Histology quality

#### 4.1.1 Staining intensity

Since we had previously found slight differences in the background staining intensity of mice brains fixed with SSS,^9^ we assessed intensity of the background staining on a scale of pale to dark as it could impact the visibility of the landmarks on the histology sections, consequently affecting the registration quality. We found a higher frequency of dark sections (which indicates the best quality) in the blocks fixed with the three solutions, reflecting no impact of the chemical composition of the different solutions on the staining background when considered altogether. However, the staining intensity seems to be affected by the staining type. Indeed, HC showed significantly more dark sections, which was expected as our previous assessments had also shown strong labeling and no impact of fixative.^10^ Furthermore, antigenicity preservation (and therefore staining intensity) was not homogeneous between the antibodies and the fixatives.^10^ Our results showed that IHC without AR have significantly more sections with both pale and dark backgrounds, while IHC with AR showed more heterogeneous sections. This could be caused by the fact that AR increases antigen staining but also decreases non-specific background labeling.

#### 4.1.2 Gray matter to white matter contrast

Gray to white matter contrast was also assessed since it could have an impact on the chosen landmarks and therefore on the registration quality. We found that SSS-fixed brain blocks showed a higher number of sections with blurred contrast compared to the other solutions. This was consistent with our previous observations showing a slightly lower contrast of the MRIs in SSS-fixed brains.^12^ This is also consistent with previous findings in histochemical sections where SSS-fixed brains showed paler neurons in Cresyl violet stain, heterogeneous fibers in Luxol fast blue, and lower quality in Bielschowsky’s staining.^10^ GM-WM contrast was also affected by the staining type, where IHC without AR showed more sections without contrast. Again, this is because AR increases the labeling of the cells and therefore shows a higher differentiation between gray and white matter layers, otherwise this is lacking when no antigen retrieval is applied to the procedure.

### 4.2 Registration quality

The number of tags that were used to manually register the T2-TurboRARE sequence with histology sections was significantly different in the brains fixed with the three solutions. AFS and NBF-fixed blocks required less tags than the SSS-fixed blocks, reflecting a more difficult registration with the latter. This is consistent with our previous findings that SSS-fixed tissue is more friable and difficult to manipulate than the other fixatives, resulting in more tears and folding of the tissue,^10^ and therefore more distortions compared to the MRI sections. This is also supported by our quantitative analysis of the percentage of Good quality tissue masks, where we found that SSS-fixed blocks showed less overlap with the tissue block masks than AFS-fixed sections, and that a higher percentage of Torn tissue was found in the SSS-fixed brains.

Regarding the relation between registration quality and the staining types, we found no difference in the number of tags required to initiate manual alignment between the three solutions. However, we found a higher Good tissue quality overlap and less Folded and Torn tissue in the histology sections that were processed with IHC without an antigen retrieval protocol. Indeed, we expected that it would be more difficult to register the sections when an AR protocol was conducted prior to IHC since it is known to enhance tissue tearing and tissue loss.^10,35^ However, AR staining being darker (higher contrast with the cells which resulted in a higher GM-WM contrast), it was unexpectedly easy to put the tags in most cases. Therefore, we suggest using an AR protocol in IHC even if this results in a more difficult manipulation of the sections, since it also results in a better staining and therefore, better correlates with MRI findings.

### 4.3 Groups of comparison

#### 4.3.1 Fixatives

Immersion of freshly extracted brains in 10% NBF is a ubiquitous fixation protocol in brain banks.^8,34^ This is currently coined as the control group since neuroscientific findings have been made with these brains so far. However, NBF solution is not ideal for anatomy dissection and also show inconveniences in histology and MRIs since the surface of the brain shows signs of over-fixation, while the deep white matter and nucleus is more poorly preserved.^36,37^ Furthermore, brain banks offer very few and small tissue samples to researchers, so we suggest that brains from anatomy laboratories may be used as an alternative. Moreover, perfusion fixation with NBF is feasible,^38^ and can be combined with perfusing the rest of the body with another solution.^39^ This would allow the adequate preparation of cadaveric specimens for gross dissection and surgical training along with optimal brain preservation for histochemical and immunohistochemical processing.

However, practices in multiple anatomy laboratories are not yet ideal and use alcohol-formaldehyde solutions and salt-saturated solution as alternatives.

Our results support that the alcohol-formaldehyde solution provides a good alternative for fixation, since it preserves brain MRI contrasts,^11,12^ histology quality and manual registration of both modalities in a good manner, as well as for combining gross anatomy dissection and histology assessment of different tissues.^41-48^

Conversely, SSS-fixed brains are of lesser quality, impacting the registration of MRI and histology images, although still feasible and of sufficient quality. These brain blocks require greater care in manipulation and histology protocols to achieve registration results closer to those obtained with the other two solutions. This is probably due to the lower amount of formaldehyde in its formulation,^40^ leading to a poorer preservation of the tissue integrity, as explained in our previous findings.^10^

#### 4.3.2 Staining type

IHC without AR group included the 4 labeled sections per specimen (NeuN, GFAP, Iba1 and PLP) that did not go through an AR procedure, while IHC with AR did. We decided to separate these two groups since we considered that heat-induced antigen retrieval could impact: 1-GM-WM contrast and staining intensity and 2-tears and folding of the tissue as we observed in a previous study of our group.^10^ Furthermore, we chose to include 4 IHC stains (NeuN, GFAP, Iba1, and PLP) to label the main brain cell populations (i.e., neurons, astrocytes, microglia and myelin), which are commonly used in histopathology diagnosis, using protocols already validated in our previous study, which made this new assessment more robust.^10^

The HC protocols included the two levels of sections using five different stains (i.e., ten slices per specimen), to label: 1-the endoplasmic reticulum and cellular nuclei, which stained cell bodies of neurons and glial cells, using a Cresyl Violet and a Neutral red staining^49^; 2-the myelin of oligodendrocytes using Luxol Fast Blue^50^, frequently used in multiple sclerosis, Alzheimer’s disease or cerebrovascular diseases^17,24,26-28,30-32,51-53^; 3-the ferric iron aggregation in the neuropil of the brain, using a Prussian blue staining (Perl’s stain),^54-56^ commonly used in different white matter pathologies detected by MRI, or again, neurodegenerative diseases^28,57-61^; 4-the nerve fibers and neurofibrils using Bielschowsky’s silver staining method,^62^ used typically to demonstrate axonal tangles and senile plaques,^63-70^ and 5-the cell bodies and small capillaries using H&E staining, the most common histopathological stain used in diagnosis of neuropathology.^26,27,29,51,58,69,71-73^ HC stains seem generally more robust than IHC, giving the highest staining intensity and sharpest GM-WM contrast. This can be related to the fact that antigens and epitopes recognition, which can be modified by cross-linking of the proteins related to chemical fixation, is not involved in these HC staining protocols.

### 4.4 Limitations

We acknowledge that this study was conducted with a small convenience sample. The small sample size resulted in heterogeneous groups of comparison regarding the staining types. More sections were processed with HC (10 per specimen) than IHC (4 with AR and 4 without AR per specimen); it is therefore not possible to know the impact of each stain (5 HC + 8 IHC) considered individually, nor that of individual antigens. Another limitation of this study is that our blocks were too large to be cut as they were scanned (we had to trim them to fit them in the vibratome). This resulted in an impossible direct comparison on tissue loss and colocalization between the histology and MRIs. However, we tried to overcome this issue by segmenting masks of the photographs of the tissue blocks after being trimmed, of the MRIs, and of the histology to facilitate the comparisons. Finally, we also had to deal with sections that showed Good histology results but that were off plane (Label 4 of the masks), which resulted in poor registration quality (high number of tags) since they were not cut parallel to the MRI scans. However, this is not a problem related to the fixative groups or the staining type, but it could have impacted the overall results. This problem could be resolved in future studies by applying an automatic nonlinear registration to the histology-MRI manual registration (Note: our in-house pipelines are still under development).

### 4.4 Conclusion

We conclude that brain blocks fixed with solutions used in gross anatomy laboratories, preferably an AFS, may be used for registration of histology photomicrographs and high-resolution MRIs of the blocks, allowing to take advantage of the two most common neuroimaging modalities. This is promising for neuroscientists interested in using larger brain samples from anatomy laboratories to correlate histology findings with MRI and possibly increase knowledge on biomarkers of normal aging or neurodegenerative disease in *post-mortem* diagnosis.

## Data availability statement

We have used the Minc ToolKit developed by the Brain Imaging Center of the Montreal Neurological Institute (McGill University) which is publicly available at: https://bic-mni.github.io.

## Acknowledgements

We would like to thank the generosity of the body donors and their families for making our projects possible. We also acknowledge funding resources and the anatomy laboratory staff of the Université du Québec à Trois-Rivières for their support and help at the lab. We would also like to thank the Cerebral Imaging Center staff (at the microscopy platform and Roqaie Moqadam) who helped in tissue management.

## References

1. Feczko E, Augustinack JC, Fischl B, Dickerson BC. An MRI-based method for measuring volume, thickness and surface area of entorhinal, perirhinal, and posterior parahippocampal cortex. Neurobiol Aging. Mar 2009;30(3):420–31. doi:10.1016/j.neurobiolaging.2007.07.023

2. Apostolova LG, Zarow C, Biado K, et al. Relationship between hippocampal atrophy and neuropathology markers: a 7T MRI validation study of the EADC-ADNI Harmonized Hippocampal Segmentation Protocol. Alzheimers Dement. Feb 2015;11(2):139–50. doi:10.1016/j.jalz.2015.01.001

3. Meyer CR, Moffat BA, Kuszpit KK, et al. A methodology for registration of a histological slide and in vivo MRI volume based on optimizing mutual information. Mol Imaging. Jan-Mar 2006;5(1):16–23.

4. Dadar M, Mahmoud S, Zhernovaia M, Camicioli R, Maranzano J, Duchesne S. White matter hyperintensity distribution differences in aging and neurodegenerative disease cohorts. Neuroimage Clin. 2022;36:103204. doi:10.1016/j.nicl.2022.103204

5. Cardoso JR, Pereira LM, Iversen MD, Ramos AL. What is gold standard and what is ground truth? Dental Press J Orthod. Sep-Oct 2014;19(5):27–30. doi:10.1590/2176-9451.19.5.027-030.ebo

6. Hoffmann A, Bredno J, Wendland MF, et al. Validation of in vivo magnetic resonance imaging blood-brain barrier permeability measurements by comparison with gold standard histology. Stroke. Jul 2011;42(7):2054–60. doi:10.1161/strokeaha.110.597997

7. Carlos AF, Poloni TE, Medici V, Chikhladze M, Guaita A, Ceroni M. From brain collections to modern brain banks: A historical perspective. Alzheimers Dement (N Y). 2019;5:52–60. doi:10.1016/j.trci.2018.12.002

8. Vonsattel JP, Amaya Mdel P, Cortes EP, Mancevska K, Keller CE. Twenty-first century brain banking: practical prerequisites and lessons from the past: the experience of New York Brain Bank, Taub Institute, Columbia University. Cell Tissue Bank. Sep 2008;9(3):247–58. doi:10.1007/s10561-008-9079-y

9. Frigon E-M, Dadar M, Boire D, Maranzano J. Antigenicity is preserved with fixative solutions used in human gross anatomy: A mice brain immunohistochemistry study. Original Research. Front Neuroanat. 2022-October-12 2022;16 doi:10.3389/fnana.2022.957358

10. Frigon EM, Gérin-Lajoie A, Dadar M, Boire D, Maranzano J. Comparison of histological procedures and antigenicity of human post-mortem brains fixed with solutions used in gross anatomy laboratories. Front Neuroanat. 2024;18:1372953. doi:10.3389/fnana.2024.1372953

11. Frigon EM, Pharand P, Gérin-Lajoie A, et al. T1- and T2-weighted MRI signal and histology findings in suboptimally fixed human brains. J Neurosci Methods. Dec 2024;412:110301. doi:10.1016/j.jneumeth.2024.110301

12. Maranzano J, Dadar M, Bertrand-Grenier A, et al. A novel ex vivo, in situ method to study the human brain through MRI and histology. J Neurosci Methods. Nov 1 2020;345:108903. doi:10.1016/j.jneumeth.2020.108903

13. Choe AS, Gao Y, Li X, Compton KB, Stepniewska I, Anderson AW. Accuracy of image registration between MRI and light microscopy in the ex vivo brain. Magnetic Resonance Imaging. 2011/06/01/ 2011;29(5):683–692. doi:10.1016/j.mri.2011.02.022

14. Goubran M, de Ribaupierre S, Hammond RR, et al. Registration of in-vivo to ex-vivo MRI of surgically resected specimens: a pipeline for histology to in-vivo registration. J Neurosci Methods. Feb 15 2015;241:53–65. doi:10.1016/j.jneumeth.2014.12.005

15. Singh M, Rajagopalan A, Kim TS, et al. Co-registration of In-Vivo Human MRI Brain Images to Postmortem Histological Microscopic Images. Int J Imaging Syst Technol. 2008;18(5-6):325–335. doi:10.1002/ima.v18:5/6

16. Bobinski M, de Leon MJ, Wegiel J, et al. The histological validation of post mortem magnetic resonance imaging-determined hippocampal volume in Alzheimer’s disease. Neuroscience. 2000;95(3):721–5. doi:10.1016/s0306-4522(99)00476-5

17. Bronge L, Bogdanovic N, Wahlund LO. Postmortem MRI and histopathology of white matter changes in Alzheimer brains. A quantitative, comparative study. Dement Geriatr Cogn Disord. 2002;13(4):205–12. doi:10.1159/000057698

18. Frigerio I, Boon BDC, Lin CP, et al. Amyloid-β, p-tau and reactive microglia are pathological correlates of MRI cortical atrophy in Alzheimer’s disease. Brain Commun. 2021;3(4):fcab281. doi:10.1093/braincomms/fcab281

19. Goubran M, Crukley C, de Ribaupierre S, Peters TM, Khan AR. Image registration of ex-vivo MRI to sparsely sectioned histology of hippocampal and neocortical temporal lobe specimens. Neuroimage. Dec 2013;83:770–81. doi:10.1016/j.neuroimage.2013.07.053

20. Mega MS, Chen SS, Thompson PM, et al. Mapping histology to metabolism: coregistration of stained whole-brain sections to premortem PET in Alzheimer’s disease. Neuroimage. Feb 1997;5(2):147–53. doi:10.1006/nimg.1996.0255

21. Scheltens P, Barkhof F, Leys D, Wolters EC, Ravid R, Kamphorst W. Histopathologic correlates of white matter changes on MRI in Alzheimer’s disease and normal aging. Neurology. May 1995;45(5):883–8. doi:10.1212/wnl.45.5.883

22. Pozorski V, Oh JM, Adluru N, et al. Longitudinal white matter microstructural change in Parkinson’s disease. Hum Brain Mapp. Oct 2018;39(10):4150–4161. doi:10.1002/hbm.24239

23. Stille M, Smith EJ, Crum WR, Modo M. 3D reconstruction of 2D fluorescence histology images and registration with in vivo MR images: application in a rodent stroke model. J Neurosci Methods. Sep 30 2013;219(1):27–40. doi:10.1016/j.jneumeth.2013.06.003

24. Gouw AA, Seewann A, van der Flier WM, et al. Heterogeneity of small vessel disease: a systematic review of MRI and histopathology correlations. J Neurol Neurosurg Psychiatry. Feb 2011;82(2):126–35. doi:10.1136/jnnp.2009.204685

25. Grinberg LT, Amaro Junior E, da Silva AV, et al. Improved detection of incipient vascular changes by a biotechnological platform combining post mortem MRI in situ with neuropathology. J Neurol Sci. Aug 15 2009;283(1-2):2–8. doi:10.1016/j.jns.2009.02.327

26. Humphreys CA, Jansen MA, Muñoz Maniega S, et al. A protocol for precise comparisons of small vessel disease lesions between ex vivo magnetic resonance imaging and histopathology. Int J Stroke. Apr 2019;14(3):310–320. doi:10.1177/1747493018799962

27. Lahna D, Roese N, Woltjer R, et al. Postmortem 7T MRI for guided histopathology and evaluation of cerebrovascular disease. J Neuropathol Exp Neurol. Dec 19 2022;82(1):57–70. doi:10.1093/jnen/nlac103

28. Roseborough AD, Langdon KD, Hammond R, et al. Post-mortem 7 Tesla MRI detection of white matter hyperintensities: A multidisciplinary voxel-wise comparison of imaging and histological correlates. NeuroImage: Clinical. 2020/01/01/ 2020;27:102340. doi:10.1016/j.nicl.2020.102340

29. Solé-Guardia G, Custers E, de Lange A, et al. Association between hypertension and neurovascular inflammation in both normal-appearing white matter and white matter hyperintensities. Acta Neuropathol Commun. Jan 4 2023;11(1):2. doi:10.1186/s40478-022-01497-3

30. Filippi M, Brück W, Chard D, et al. Association between pathological and MRI findings in multiple sclerosis. Lancet Neurol. Feb 2019;18(2):198–210. doi:10.1016/s1474-4422(18)30451-4

31. Laule C, Kozlowski P, Leung E, Li DK, Mackay AL, Moore GR. Myelin water imaging of multiple sclerosis at 7 T: correlations with histopathology. Neuroimage. May 1 2008;40(4):1575–80. doi:10.1016/j.neuroimage.2007.12.008

32. Laule C, Moore GRW. Myelin water imaging to detect demyelination and remyelination and its validation in pathology. Brain Pathol. Sep 2018;28(5):750–764. doi:10.1111/bpa.12645

33. Moll NM, Rietsch AM, Thomas S, et al. Multiple sclerosis normal-appearing white matter: pathology-imaging correlations. Ann Neurol. Nov 2011;70(5):764–73. doi:10.1002/ana.22521

34. Dadar M, Sanches LG, Fouquet J, et al. The Douglas-Bell Canada Brain Bank Post-mortem Brain Imaging Protocol. Apuerture Neuro. 2024;4 doi:10.52294/001c.123347

35. Evers P, Uylings HB, Suurmeijer AJ. Antigen retrieval in formaldehyde-fixed human brain tissue. Methods. Jun 1998;15(2):133–40. doi:10.1006/meth.1998.0616

36. Garman RH. Artifacts in routinely immersion fixed nervous tissue. Toxicol Pathol. 1990;18(1 Pt 2):149–53. doi:10.1177/019262339001800120

37. Beach TG, Tago H, Nagai T, Kimura H, McGeer PL, McGeer EG. Perfusion-fixation of the human brain for immunohistochemistry: comparison with immersion-fixation. J Neurosci Methods. Mar 1987;19(3):183–92. doi:10.1016/s0165-0270(87)80001-8

38. McFadden WC, Walsh H, Richter F, et al. Perfusion fixation in brain banking: a systematic review. Acta Neuropathol Commun. Sep 5 2019;7(1):146. doi:10.1186/s40478-019-0799-y

39. Insausti R, Insausti AM, Muñoz López M, et al. Ex vivo, in situ perfusion protocol for human brain fixation compatible with microscopy, MRI techniques, and anatomical studies. Methods. Front Neuroanat. 2023-March-23 2023;17 doi:10.3389/fnana.2023.1149674

40. Coleman R, Kogan I. An improved low-formaldehyde embalming fluid to preserve cadavers for anatomy teaching. J Anat. Apr 1998;192 (Pt 3)(Pt 3):443–6. doi:10.1046/j.1469-7580.1998.19230443.x

41. Venne G, Zec ML, Welte L, Noel G. Qualitative and quantitative comparison of Thiel and phenol-based soft-embalmed cadavers for surgery training. Anat Histol Embryol. May 2020;49(3):372–381. doi:10.1111/ahe.12539

42. Tomalty D, Pang SC, Ellis RE. Preservation of neural tissue with a formaldehyde-free phenol-based embalming protocol. Clin Anat. Mar 2019;32(2):224–230. doi:10.1002/ca.23290

43. Rahman SMN, Alam T, Alam NN. Preservation of Histology by Phenol-Based Fixative: Mini Review of Recent Findings %J International Journal of Morphology. 2021;39:50–56.

44. O’Sullivan E, Mitchell BS. An improved composition for embalming fluid to preserve cadavers for anatomy teaching in the United Kingdom. J Anat. Apr 1993;182 (Pt 2)(Pt 2):295–7.

45. Martins-Costa C, Nunes TC, Anjos-Ramos LD. Anatomo-comparative study of formaldehyde, alcohol, and saturated salt solution as fixatives in Wistar rat brains. Anat Histol Embryol. Nov 2022;51(6):740–745. doi:10.1111/ahe.12852

46. Hopwood D, Slidders W, Yeaman GR. Tissue fixation with phenol-formaldehyde for routine histopathology. Histochem J. Apr 1989;21(4):228–34. doi:10.1007/bf01747525

47. Benet A, Rincon-Torroella J, Lawton MT, González Sánchez JJ. Novel embalming solution for neurosurgical simulation in cadavers. J Neurosurg. May 2014;120(5):1229–37. doi:10.3171/2014.1.Jns131857

48. Barton DP, Davies DC, Mahadevan V, et al. Dissection of soft-preserved cadavers in the training of gynaecological oncologists: report of the first UK workshop. Gynecol Oncol. Jun 2009;113(3):352–6. doi:10.1016/j.ygyno.2009.02.012

49. Eilam-Altstädter R, Las L, Witter MP, Ulanovsky N. Methods. In: Eilam-Altstädter R, Las L, Witter MP, Ulanovsky N, eds. Stereotaxic Brain Atlas of the Egyptian Fruit Bat. Academic Press; 2021:1–9.

50. Lindberg MR, Lamps LW. Central Nervous System. In: Lindberg MR, Lamps LW, eds. Diagnostic Pathology: Normal Histology (Second Edition). Elsevier; 2018:108–111.

51. Li K, Rashid T, Li J, et al. Postmortem Brain Imaging in Alzheimer’s Disease and Related Dementias: The South Texas Alzheimer’s Disease Research Center Repository. J Alzheimers Dis. 2023;96(3):1267–1283. doi:10.3233/jad-230389

52. Kuhlmann T, Ludwin S, Prat A, Antel J, Brück W, Lassmann H. An updated histological classification system for multiple sclerosis lesions. Acta Neuropathol. Jan 2017;133(1):13–24. doi:10.1007/s00401-016-1653-y

53. Ouellette R, Mangeat G, Polyak I, et al. Validation of Rapid Magnetic Resonance Myelin Imaging in Multiple Sclerosis. Ann Neurol. May 2020;87(5):710–724. doi:10.1002/ana.25705

54. Churukian CJ. 14 - Pigments and Minerals. In: Bancroft JD, Gamble M, eds. Theory and Practice of Histological Techniques (Sixth Edition). Churchill Livingstone; 2008:233–259.

55. Guindi M. 11 - Liver Disease in Iron Overload. In: Saxena R, ed. Practical Hepatic Pathology: a Diagnostic Approach (Second Edition). Elsevier; 2018:151–165.

56. Xiao G, Li H, Zhao M, Zhou B. Chapter Eight - Assessing metal ion transporting activity of ZIPs: Intracellular zinc and iron detection. In: Hu J, ed. Methods in Enzymology. Academic Press; 2023:157–184.

57. Connor JR, Benkovic SA. Iron regulation in the brain: histochemical, biochemical, and molecular considerations. Ann Neurol. 1992;32 Suppl:S51–61. doi:10.1002/ana.410320710

58. Kruer MC. The neuropathology of neurodegeneration with brain iron accumulation. Int Rev Neurobiol. 2013;110:165–94. doi:10.1016/b978-0-12-410502-7.00009-0

59. Dusek P, Hofer T, Alexander J, Roos PM, Aaseth JO. Cerebral Iron Deposition in Neurodegeneration. Biomolecules. May 17 2022;12(5) doi:10.3390/biom12050714

60. Stüber C, Morawski M, Schäfer A, et al. Myelin and iron concentration in the human brain: a quantitative study of MRI contrast. Neuroimage. Jun 2014;93 Pt 1:95–106. doi:10.1016/j.neuroimage.2014.02.026

61. van Duijn S, Nabuurs RJ, van Duinen SG, Natté R. Comparison of histological techniques to visualize iron in paraffin-embedded brain tissue of patients with Alzheimer’s disease. J Histochem Cytochem. Nov 2013;61(11):785–92. doi:10.1369/0022155413501325

62. Ogawa Y. Silver Impregnation Method for Neurons Using Synthetic Surfactants: A Contribution to Golgi Method. In: Conn PM, ed. Quantitative and Qualitative Microscopy. Methods in Neurosciences; 1990:chap STAINING TECHNOLOGY.

63. Stahnisch FW. Max Bielschowsky (1869-1940). J Neurol. Mar 2015;262(3):792–4. doi:10.1007/s00415-014-7544-z

64. Wisniewski HM, Wen GY, Kim KS. Comparison of four staining methods on the detection of neuritic plaques. Acta Neuropathol. 1989;78(1):22–7. doi:10.1007/bf00687398

65. Love S, Nicoll JA. Comparison of modified Bielschowsky silver impregnation and anti-ubiquitin immunostaining of cortical and nigral Lewy bodies. Neuropathol Appl Neurobiol. Dec 1992;18(6):585–92. doi:10.1111/j.1365-2990.1992.tb00830.x

66. Allsop D. Introduction to Alzheimer’s disease. Methods Mol Med. 2000;32:1–21. doi:10.1385/1-59259-195-7:1

67. Switzer RC, 3rd. Application of silver degeneration stains for neurotoxicity testing. Toxicol Pathol. Jan-Feb 2000;28(1):70–83. doi:10.1177/019262330002800109

68. Uchihara T. Silver diagnosis in neuropathology: principles, practice and revised interpretation. Acta Neuropathol. May 2007;113(5):483–99. doi:10.1007/s00401-007-0200-2

69. Kovacs GG, Budka H. Current concepts of neuropathological diagnostics in practice: neurodegenerative diseases. Clin Neuropathol. Sep-Oct 2010;29(5):271–88. doi:10.5414/npp29271

70. Elobeid A, Rantakömi S, Soininen H, Alafuzoff I. Alzheimer’s disease-related plaques in nondemented subjects. Alzheimers Dement. Sep 2014;10(5):522–9. doi:10.1016/j.jalz.2012.12.009

71. Vakrakou AG, Brinia ME, Svolaki I, Argyrakos T, Stefanis L, Kilidireas C. Immunopathology of Tumefactive Demyelinating Lesions-From Idiopathic to Drug-Related Cases. Front Neurol. 2022;13:868525. doi:10.3389/fneur.2022.868525

72. Beach TG, Sue LI, Scott S, et al. Cerebral white matter rarefaction has both neurodegenerative and vascular causes and may primarily be a distal axonopathy. J Neuropathol Exp Neurol. May 25 2023;82(6):457–466. doi:10.1093/jnen/nlad026

73. Fischer AH, Jacobson KA, Rose J, Zeller R. Hematoxylin and eosin staining of tissue and cell sections. CSH Protoc. May 1 2008;2008:pdb.prot4986. doi:10.1101/pdb.prot4986

